# Food intake habits and preference affects the activity of medial frontal cortex during eating; functional near-infrared spectroscopy is potential biosensor for appetite study

**DOI:** 10.64898/2026.02.09.704900

**Authors:** Yusuke Takatsuru, Yuka Sekine, Hideyasu Sato, Tomoko Osera

## Abstract

Even if they have no dementia, some elderly people find it difficult to imagine the food they may want to eat. However, research into treatments for anorexia in elderly people has not progressed sufficiently due to the lack of a method that can easily measure brain function in clinical settings. In this study, we aim to clarify the relationship of food-dependent medial prefrontal cortex (MPFC) activity with food preference and intake frequency by functional near-infrared spectroscopy (fNIRS) to determine ways to treat the appetite loss and have difficulty in explain what they want to eat. For this purpose, we firstly establish the methodology using young participants experiment. All young participants were asked about their food preferences and intake frequency using a questionnaire, and they were instructed to look and then eat the control dish (CD: typical Japanese home-cooked meal) and their preferred dish (PD: each participant purchased the dish themselves on the day of the experiments) on separate days, and activity of MPFC in each participant was recorded by fNIRS. We found that activity of MPFC during “just-looking” and “eating” were affected by food intake habits and preference. Especially, activity of MPFC during CD eating was affected by food preference (has dislike food or not). We concluded that the activity of MPFC during eating of dishes varies depending on the food intake habits and fNIRS could be a potential technique for estimating such activity.

## Introduction

With the increasing number of elderly people in Japan (the percentage of the elderly has increased to 29.1%)[1], we are facing the problem of appetite loss among them. Chewing problems, depressive symptoms, and polypharmacy among others decrease the appetite of the elderly [2–5]. Even if they have no dementia, some elderly people find it difficult to imagine the food they may want to eat. In the case of the elderly with swallowing problem, they “don’t want to eat” because they can only eat food paste. Morley and Silver [6] categorized the appetite loss in the elderly into; decreased demand, decreased hedonic qualities, decreased feeding drive, and increased activity of satiety factors. Appetite loss aggravates malnutrition disorders such as frailty in the elderly [7,8]. However, studies of appetite loss in the elderly are still insufficient [4]. One reason for this may be that research into the role of the brain in appetite and related food choices by easily measuring brain function in clinical settings has not progressed.

Brain activities during eating have been assessed using neuroimaging techniques such as functional magnetic resonance imaging (fMRI), positron emission tomography, and magnetoencephalography [9]. These techniques can accurately show the localization of brain activities in response to taste stimulation. However, they require their head to be hold completely stationary, which is far from the usual condition during eating. From this perspective, functional near-infrared spectroscopy (fNIRS) is one of the noninvasive techniques for detecting brain activities similarly to fMRI. Although fNIRS has a poorer spatial resolution than fMRI [10], it has a higher temporal resolution than fMRI, and a subject can move more freely and perform a wider range of tasks such as eating [11,12]. As previously reported, the activity of MPFC area 10 during the task of eating food, such as preferred and nonpreferred taste, has been studied well by fNIRS [9]. Moreover, the fNIRS setup is easy to move around compared with the MRI setup and thus, we can record signals even if participants cannot come to a special area.

We eat provided food (in our own homes, in hospitals, or schools; thus we cannot choose) or purchased food (we can choose what we want eat from shops or restaurants). What happens when we can choose the food and decide what to eat? The information of vision, olfaction, and taste affects eating behavior [13]. Such information is first processed at visual cortex, olfactory cortex, and frontal operculum/insula. Then, the amygdala, orbitofrontal cortex, and pregenual cingulate cortex process the information as a reward/affective value for food/eating. These brain areas also moderated by the lateral prefrontal cortex and hunger-related neurons. Finaly, area 10 of the medial prefrontal cortex (MPFC; decision-making), the cingulate cortex (for action output), the striatum (for habit), and the lateral hypothalamus (for autonomic and endocrine responses) control the eating behavior and related actions. Eating behavior is, thus, very complicated and affected by several factors.

Aim of this study is to clarify the relationship of the activity of MPFC with food intake by fNIRS to find satisfactory methods of mechanism of food eating decision-making in the elderly who have lost their appetite. In this study, however, we performed the experiment with young participants because of establishing the methodology. The activity of MPFC be recorded by fNIRS even if the participants cannot speak or write theirs answers to questionnaires. Thus, if we can find some correlation between the activity of MPFC and questionnaire responses for categorizing the participants, we can determine their food intake behavior/preference by using only fNIRS data, which may be useful for the elderly people especially for those having trouble in saying/writing what they want to eat. For this purpose, we prepared two types of dish. The control dish (CD: typical Japanese home-cooked meal) provided in institutions such as hospitals) and their preferred dish (PD: each participant purchased the dish themselves on the day of the experiments). We also examined the relationship of food eating decision-making with food preference, frequency of food intake, and knowledge of food. These factors could affect the decision making of eating. Some of them are correlated with different brain regions [9] and are difficult to observe by fNIRS. However, if we find some relationship of these factors with MPFC activity, it may be useful for clarifying the appetite of the elderly. Because this is a pilot study to establish the methods before applying them to elderly participants, this study was performed on young participants.

## Materials and Methods

### Procedure

The study was approved by the ethics committee of Toyo University (TU2020-011-TU2021-H-099-TU2021-H-023-TU2023-K-046) and performed in accordance with the Declaration of Helsinki. The experiments were conducted on participants who applied themselves through oral or posted recruitment. After the experiments were completed, one subject self-reported a history of eating disorders and was therefore excluded from the study. All 68 participants (33 males, 21.6 ± 0.2 years old; 35 females, 21.2 ± 0.2 years old) gave informed consent before the experiments. Three female participants were left-handed and the rest were right-handed. To gather methodological knowledge, we observe healthy young participants in this study. After obtaining informed consent, all participants filled up questionnaires on food intake frequency and preference and underwent the food intake test by fNIRS recording.

### Measures

#### Food intake frequency and preference

All participants were asked about their food intake frequency, food preferences, food-related behavior, and knowledge of food using a questionnaire before starting the food intake test (Supplemental Tables 1 and 2). Their knowledge of seasonal foods was analyzed on the basis of the score of summation (one correct answer was given one point with 20 points as the maximum). Food intake frequency was determined using the Frequency Questionnaire Based on Food Groups (FFQg) (Ver. 6, Addin software of EIYO-KUN Ver. 9, Kenpakusya, Bunkyo-ku, Tokyo, Japan) [21,22] which included the information on gender and age (we only obtained the birth year. The participants were defined to have been born on January 1st, and their age was determined on the basis of the date given in the FFQg form).

#### Food intake test

All participants ate the control dish (CD: typical Japanese home-cooked meal) and their preferred dish (PD: each participant purchased the dish themselves on the day of the experiments) on separate days. All experiments were performed during lunchtime (11:00–14:00, JST). The target amount of CD was set at 34% of 1600 kcal of the daily intake for elderly women. The macronutrient balance for protein, fat, and carbohydrate (PFC) energy production was in accordance with the Dietary Reference Intakes (DRI) for Japanese [23].

CD included nine components (grains, fish, pork, soy, sugar, vegetables, sea weed, and seafood), whose adequacy was assessed in accordance with DRI (see also Appendix). The participants purchased PD themselves under the following conditions: (1) can be eaten in the university including drinks (easy to carry, highly perishable, without alcohol), (2) at amounts that can be consumed completely, and (3) the cost of food was covered by research funds. After eating the dishes, some of the participants evaluated the dishes using the visual analog scale (VAS. 0–100 mm). The participants received a compensation of 3,000-5,000 Japanese yen (which included travel fee and not excluded the cost of purchasing PD).

#### fNIRS recording

The activity of MPFC during eating was recorded using a multichannel fNIRS unit operating at wavelengths of 770 and 840 nm (OEG-16H; Spectratech Inc., Yokohama) to measure temporal changes in the concentrations of oxygenated hemoglobin (oxyHb), deoxygenated hemoglobin (deoxyHb), and total hemoglobin (totalHb) as described previously [24]. Supplemental Fig. 1 shows the position of eight infrared irradiation probes and eight infrared reception probes for the calculation of signals from the brain for information in sixteen channels. The probes were located 30 mm apart. The positions of the probes were based on the international 10–20 system used in electroencephalography and the center of the probe pad was located on Fpz (i.e., the midpoint between Fp1 and Fp2). The recording interval was 85 ms, and ten data points were averaged for smoothing, obtaining data every 0.85 s [10]. We used autocalibration in the OEG16H device before recording and if the gain was between 2000 and 100, we obtained the channel for the data analysis. If the gain was beyond this range, we reattached the probes. After reattaching and the signal gain was is detected more than our standard (more than 3 signal from cHs 7-10), we proceeded with the experiments and the data that were obtained beyond the range was omitted in the analysis. We removed the component of body movement by using the formula below [25]:

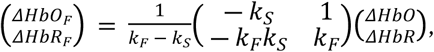

where ΔHbO_F_ is the amount of signal change for oxyhemoglobin associated with cerebral function, ΔHbD_F_ is the amount of signal change for deoxyhemoglobin associated with cerebral function, ΔHbO is the amount of signal change for oxyhemoglobin detected by fNIRS, and ΔHbD is the amount of signal change for deoxyhemoglobin detected by fNIRS. k_F_ and k_S_ are coefficients. We used 0.6 for k_F_ [25] and k_S_ was calculated for each participant using the algorithm in OEG-16H. Although this method of removing motion components may be insufficient, it is a standard feature of the device and can be easily used, which is an advantage when considering future clinical applications.

The signal data of each probe were calculated by the device using the formulas below;

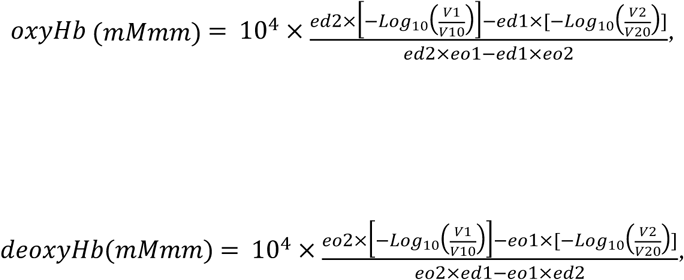

where eo1 is the molar absorption coefficient of oxyHb by 840 nm (cm^-1^/M), ed1 is the molar absorption coefficient of deoxyHb by 840 nm (cm^-1^/M), eo2 is the molar absorption coefficient of oxyHb by 770 nm (cm^-1^/M), ed2 is the molar absorption coefficient of deoxyHb by 770 nm (cm^-1^/M), V1 is the current signal evoked by 840 nm, V10 is the basal signal evoked by 840 nm, V2 is the current signal evoked by 770 nm, and V20 is the basal signal evoked by 770 nm. In our methods, eo1 (= 1022), ed1 (= 692.36), eo2 (= 650), and ed2 (= 1311.88) were automatically determined. For filtering, we use low-pass fast Fourier transform and the baseline was corrected by linear fit in device.

For each fNIRS recording, 20 s was allocated for the baseline (participants were instructed to relax and wait), 20 s for “just looking”, and a sufficient time for “eating” (the first 10 min was used for analysis). The data on oxyHb_F_ from each channel were averaged every 0.82 s, and the average signal (average oxyHb_F_) of cHs 7–10 was used as the signal from MPFC [26–31]. Hemodynamic data of fNIRS are relative values and cannot be averaged directly across channels or subjects; the z-score, as normalized data, could be averaged regardless of the unit [10]. The obtained signals were analyzed as the z-score calculated as

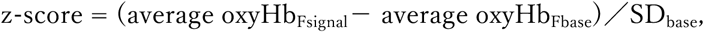

where average oxyHb_Fsignal_ is the average oxyHb_F_ of all time points, average oxyHb_Fbase_ is the average over a period of basal recording (19.68 s from baseline recording for the “just looking” condition and 19.69 s before the start of eating for the “eating” condition), and SD_base_ is the standard deviation of basal recording. When the average z-score of cH7-cH10 for 19.68 s (“just-looking”) and/or 599.65 s (“eating”) was more than 2, we considered it as a positive response and the participant with such a response was categorized as a “responder”. A CD-responder was someone who showed a positive response in MPFC during looking/eating CD. A PD-responder was someone who showed a positive response in MPFC during looking/eating PD. A both-responder was someone who showed a positive response in MPFC during looking/eating both CD and PD. A nonresponder was someone who showed a response below the threshold in MPFC during looking/eating both CD and PD.

### Statistical analyses

The obtained data were analyzed by one-way ANOVA and the Bonferroni test for post-hoc analysis, Student’s *t*-test, and Pearson correlation coefficient test for continuous variables and the Kruskal–Wallis test and Steel–Dwass test for post-hoc analysis, Mann–Whitney U-test or Fisher’s exact test using for categorical variables using the Excel analysis (BellCurve, Japan). We also performed the binary or multiple logistic regression comparison using the Excel analysis (BellCurve, Japan). Differences were considered significant at *p* < 0.05. All values are presented as mean ± SEM or median (IQR).

## Results

Differences in food preference, knowledge of seasonal foods, and the food intake frequency affect the activity of MPFC during “just-looking” and “eating” the food.

As shown in Table 1, when the participants were “just-looking” or “eating” the dishes, some of them showed a positive response in MPFC (Fig. 1). The members of the responder and nonresponder groups were different between “just-looking” and “eating”. On the basis of the four categories (CD-responders, PD-responders, both-responders, and nonresponders), we next analyzed the food intake frequency and food preference.

**Fig. 1.**
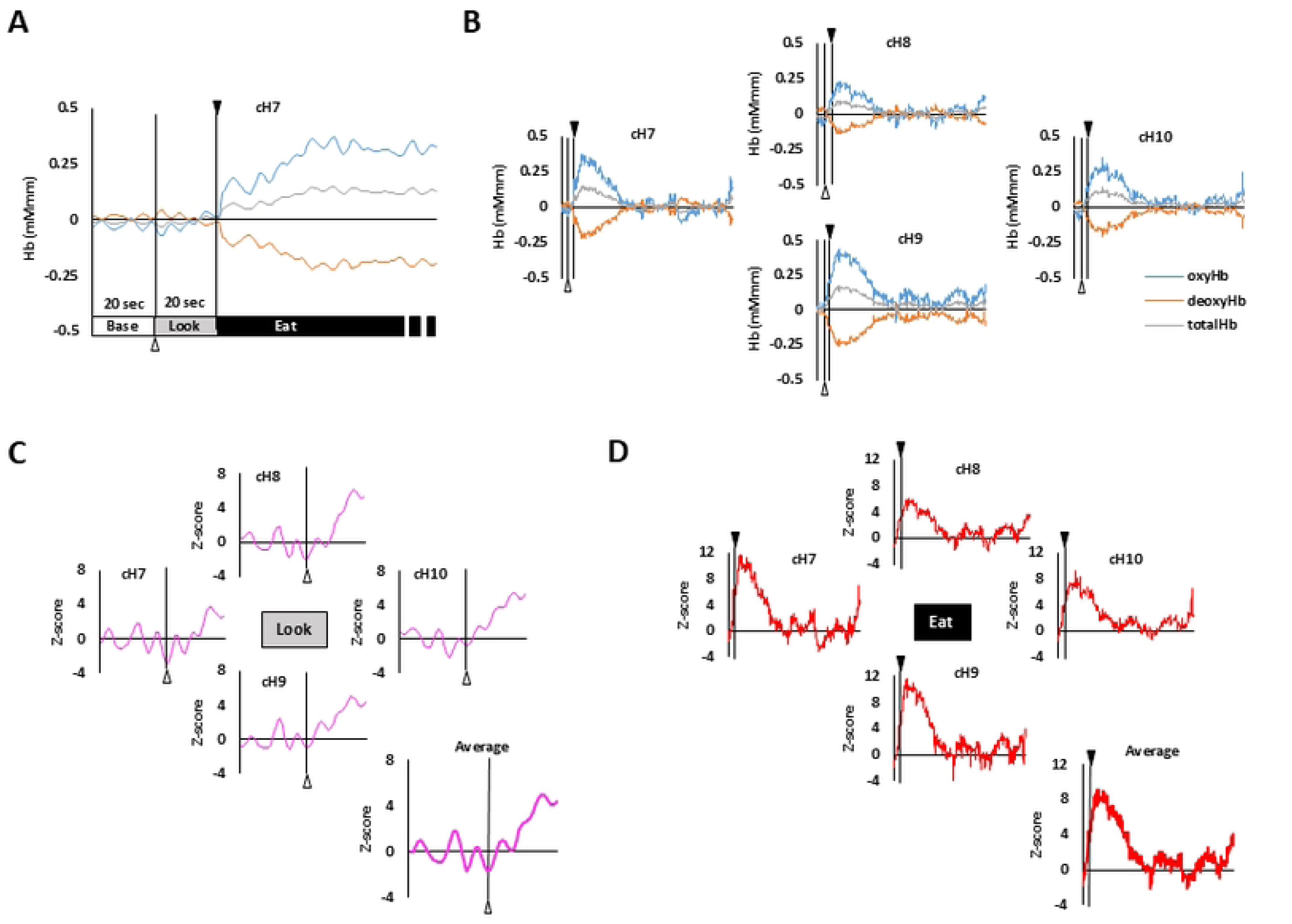
Typical signals from MPFC (A,B) and calculated z-score (C,D) in participants during eating CD. Signals from each channel were calculated as shown in Methods, and the z-score (normalized by baseline recording) is presented. In each experiment, participants were presented the food (CD or PD) after 20 s of baseline recording. After another 20 s of just looking, they started to eat. Recording was continued until they finished eating and the first 10 min of recording was used for analysis.

**Table 1.** Number of participants who showed positive responses in MPFC detected by fNIRS under each condition.

|  | CD-responders | PD-responders | Both-responders | Nonresponders |
| --- | --- | --- | --- | --- |
| “Just-looking” | 15 (4/11) | 15 (10/5) | 9 (5/4) | 29 (14/15) |
| “Eating” | 16 (8/8) | 14 (6/8) | 14 (8/6) | 24 (11/13) |

Number of participants (male/female). Responders included CD-, PD-, and both-responders.

**Table 2.** Binary logistic regression comparison of fNIRS results during “just-looking” and results of questionnaires.

|  | B(S.E.) | Odds (95% CI) | *p*-value |
| --- | --- | --- | --- |
| **CD-responders** |  |  |  |
| Eat bread at breakfast | -0.01(0.17)) | 0.99(0.71-1.38) | 0.94 |
| Eat stir frying | 0.20(0.24) | 1.22(0.77-1.94) | 0.40 |
| Oil preference | -1.11(0.57) | 0.33(0.11-1.01) | 0.053 |
| **PD-responders** |  |  |  |
| Eat bread at breakfast | 0.17(0.15) | 1.19(0.88-1.58) | 0.24 |
| Eat stir frying | -0.72(0.31) | 0.48(0.26-0.90) | **0.021** |
| Oil preference | 0.51(0.65) | 1.32(0.053-32.67) | 0.43 |
| **Both-responders** |  |  |  |
| Eat bread at breakfast | -0.031(0.21) | 0.97(0.65-1.45) | 0.88 |
| **Eat stir frying** | -0.15(0.31) | 0.86(0.47-1.58) | 0.63 |
| **Oil preference** | -1.19(0.68) | 0.31(0.081-1.15) | 0.079 |

Estimated using the Multiple logistic regression model. Gender was included as confounding factors. The inverse matrix of the second derivative matrix of the log-likelihood function on frequency of drinking protein did not exist, so it was excluded from the analysis.

As shown in supplemental Table 3, some of the factors in the questionnaires showed gender dependence. The frequency of eating carbohydrate, visit new/popular restaurants, and try the information obtained from those programs and contents, were associated with gender. However, activity of MPFC did not associate with gender.

MPFC response during “just-looking”, and food preference, knowledge of seasonal foods, and food intake frequency

As shown in Fig. 2, PD-responders during “just-looking” frequently eat bread at breakfast (CD; 1.5 ± 0.5 times/week, PD; 4.0 ± 0.8 times/week, Both; 1.4 ± 0.6 times/week, Non; 2.0 ± 0.5 times/week), drink protein (CD; 0 ± 0 times/week, PD; 1.1 ± 0.4 times/week, Both; 0.2 ± 0.2 times/week, Non; 0.1 ± 0.1 times/week). Moreover, PD-responders during “just-looking” less frequently eat stir frying (CD; 3.8 ± 0.4 times/week, PD; 2.2 ± 0.3 times/week, Both; 3.3 ± 0.4 times/week, Non; 3.5 ± 0.3 times/week) and don’t like oily taste [CD, 2(2-3), PD; 2(1-2), Both; 2(2-3), Non; 2(1-2)]. As shown in Table 2, frequency of the eat stir frying was significantly associated with the PD-responder as determined by multiple logistic regression comparison.

**Fig. 2.**
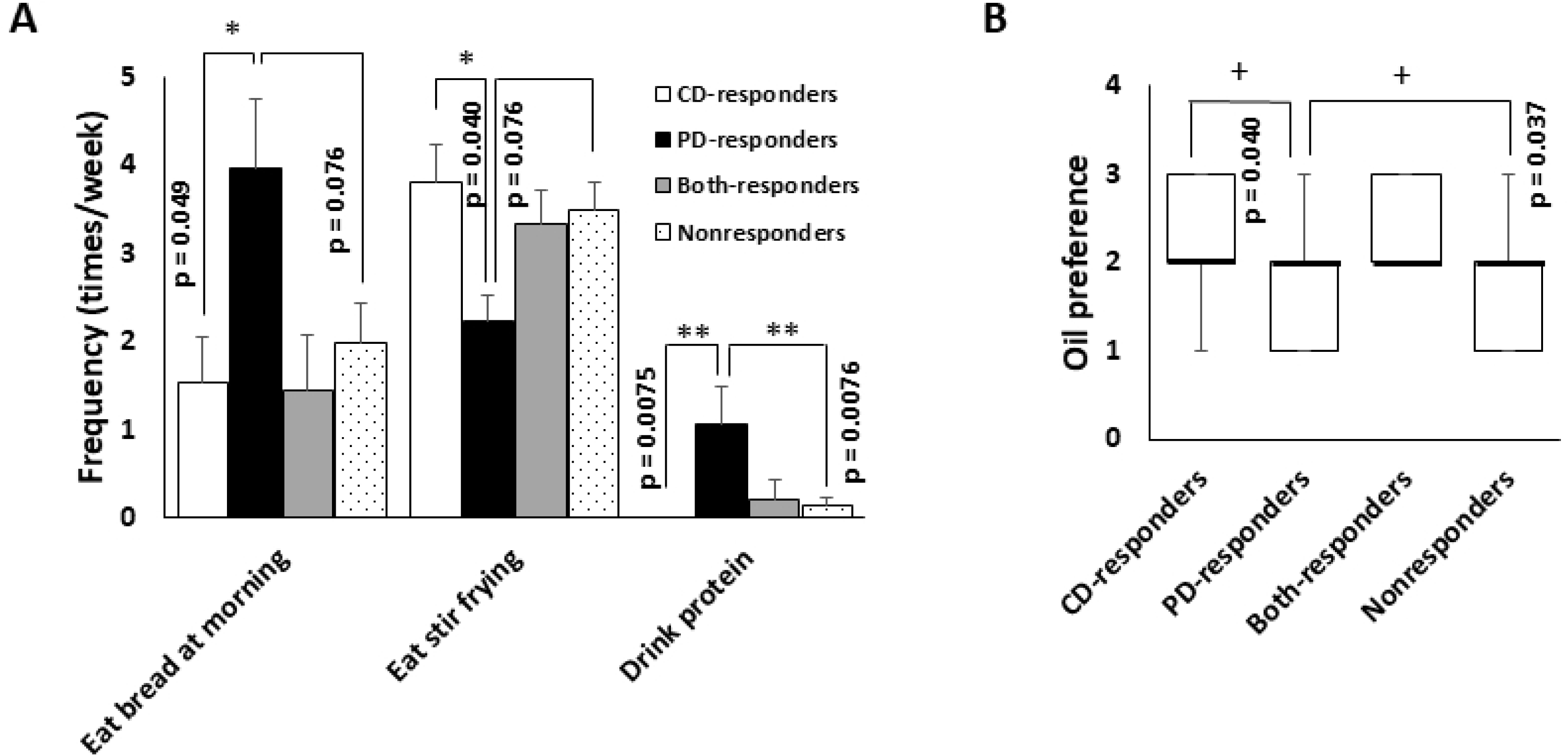
Food intake habits and preference of taste and activity of MPFC during “just-looking” the CD and PD. (A) PD-responders frequently eat bread at morning and drink protein but less frequently eat stir frying. (B) PD-responders not prefer oil. * indicates *p* < 0.05 by Bonferroni test, + indicates *p* < 0.05 by Steel–Dwass test

MPFC response during “eating”, and food preference, knowledge of seasonal foods, and food intake frequency

As shown in Fig. 3, Nonresponders tend to frequently eat soy/soy products at lunch (CD; 1.4± 0.3 times/week, PD; 0.8 ± 0.3 times/week, Both; 0.4 ± 0.2 times/week, Non; 1.9 ± 0.5 times/week). PD-responders frequently eat seaweeds (CD; 2.0 ± 0.4 times/week, PD; 3.8 ± 0.6 times/week, Both; 2.1 ± 0.6 times/week, Non; 1.8 ± 0.5 times/week). CD-responder tend to have not dislike food [CD, 0(0-1), PD; 1(1-1), Both; 1(1-1), Non; 1(0.75-1)]. As shown in Table 3, dislike food associated with the CD-responder, frequency of the eat seaweeds associated with the PD-responder, and frequency of the eat soy/soy products at lunch associated with the Both-responders as determined by multiple logistic regression comparison.

**Fig. 3.**
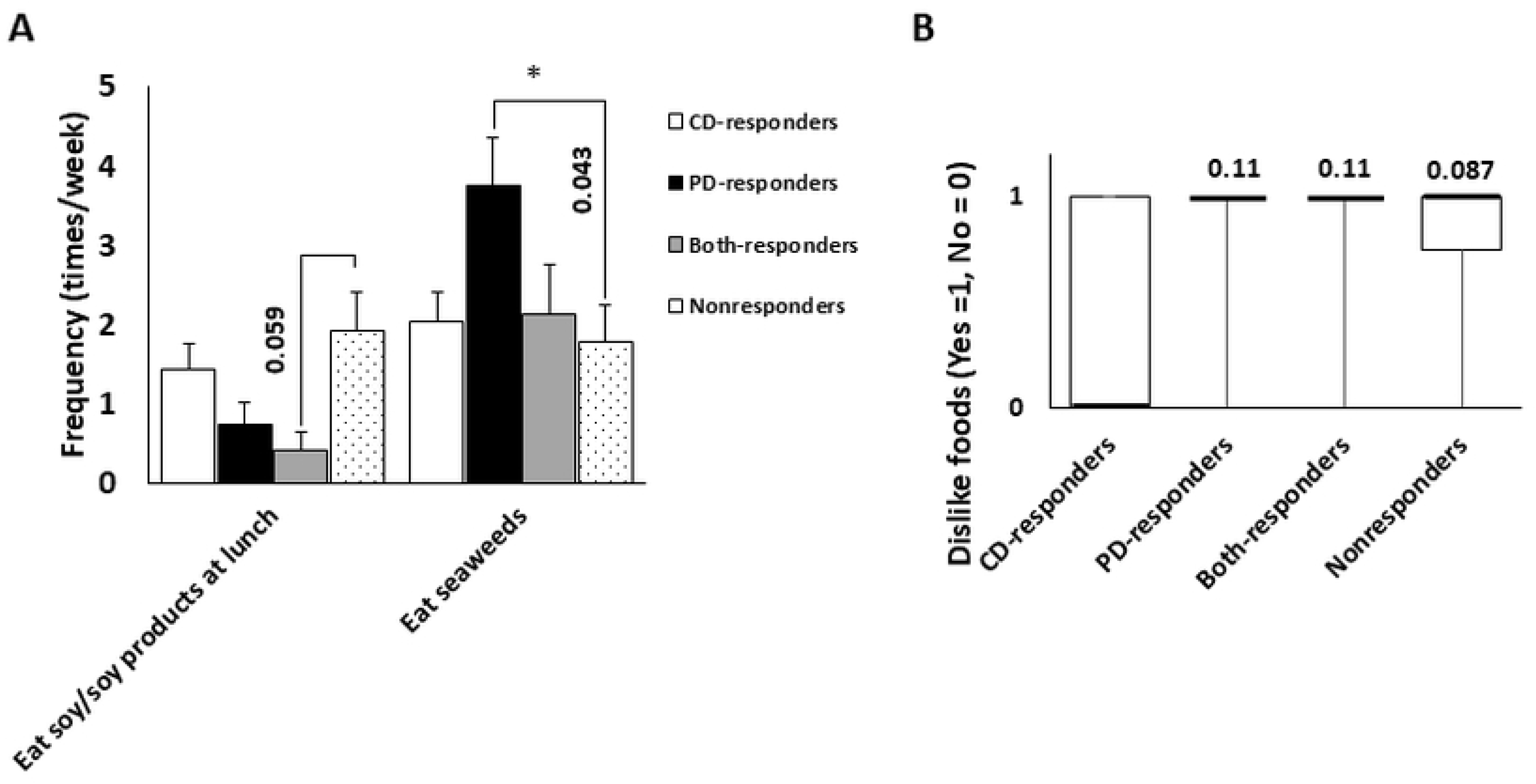
Food intake habits and preference of food and activity of MPFC during “eating” the CD and PD. (A) PD-responders frequently eat seaweeds. (B) CD-responders tends to not have dislike food.

**Table 3.** Multiple comparisons of fNIRS results during “eating” and questionnaire results.

|  | B(S.E.) | Odds (95% CI) | *p*-value |
| --- | --- | --- | --- |
| **CD-responders** |  |  |  |
| Eat soy/soy products at lunch | -0.16(0.22) | 0.85(0.56-1.30) | 0.46 |
| Eat seaweeds | 0.038(0.19) | 1.04(0.72-1.50) | 0.84 |
| Dislike food (Yes = 1, No = 0) | -1.71(0.75) | 0.18(0.041-0.79) | **0.024** |
| **PD-responders** |  |  |  |
| Eat soy/soy products at lunch | -0.63(0.35) | 0.53(0.26-1.06) | 0.073 |
| Eat seaweeds | 0.41(0.17) | 1.51(1.07-2.12) | **0.017** |
| Dislike food (Yes = 1, No = 0) | 0.12(0.91) | 1.13(0.19-6.71) | 0.89 |
| **Both-responders** |  |  |  |
| Eat soy/soy products at lunch | -1.06(0.43) | 0.35(0.15-0.81) | **0.014** |
| Eat seaweeds | 0.15(0.19) | 1.16(0.26-10.37) | 0.45 |
| **Dislike food (Yes = 1, No = 0)** | 0.50(0.19) | 1.65(0.26-10.37) | 0.59 |

Estimated using the Multiple logistic regression model. Gender was included as
confounding factors.

Activity of MPFC during “eating” and dislike food.

We next divided participants who has the dislike food or not. As shown in Table 4, number of participants who has dislike food was significantly high in female than those in male. Participants who have dislike food frequently eat meat/meat products at lunch, use better/margarine, and visit a new and popular restaurants. In contrast, participants who have no dislike food has more knowledge of the seasonal foods. As shown in Table 4, gender and the knowledge of the seasonal foods significantly associated with having the dislike food or not as determined by logistic regression comparison.

**Table 4.**
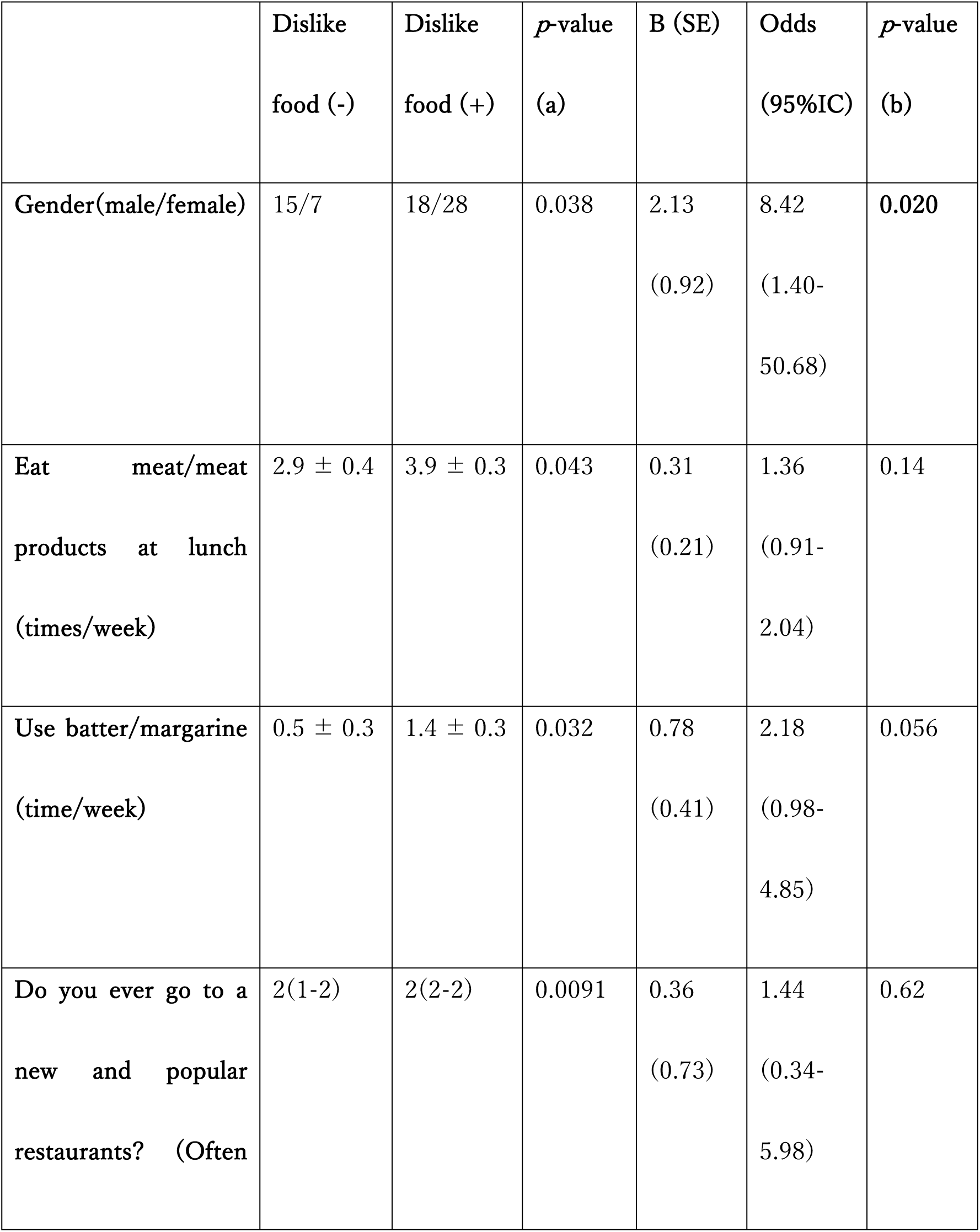
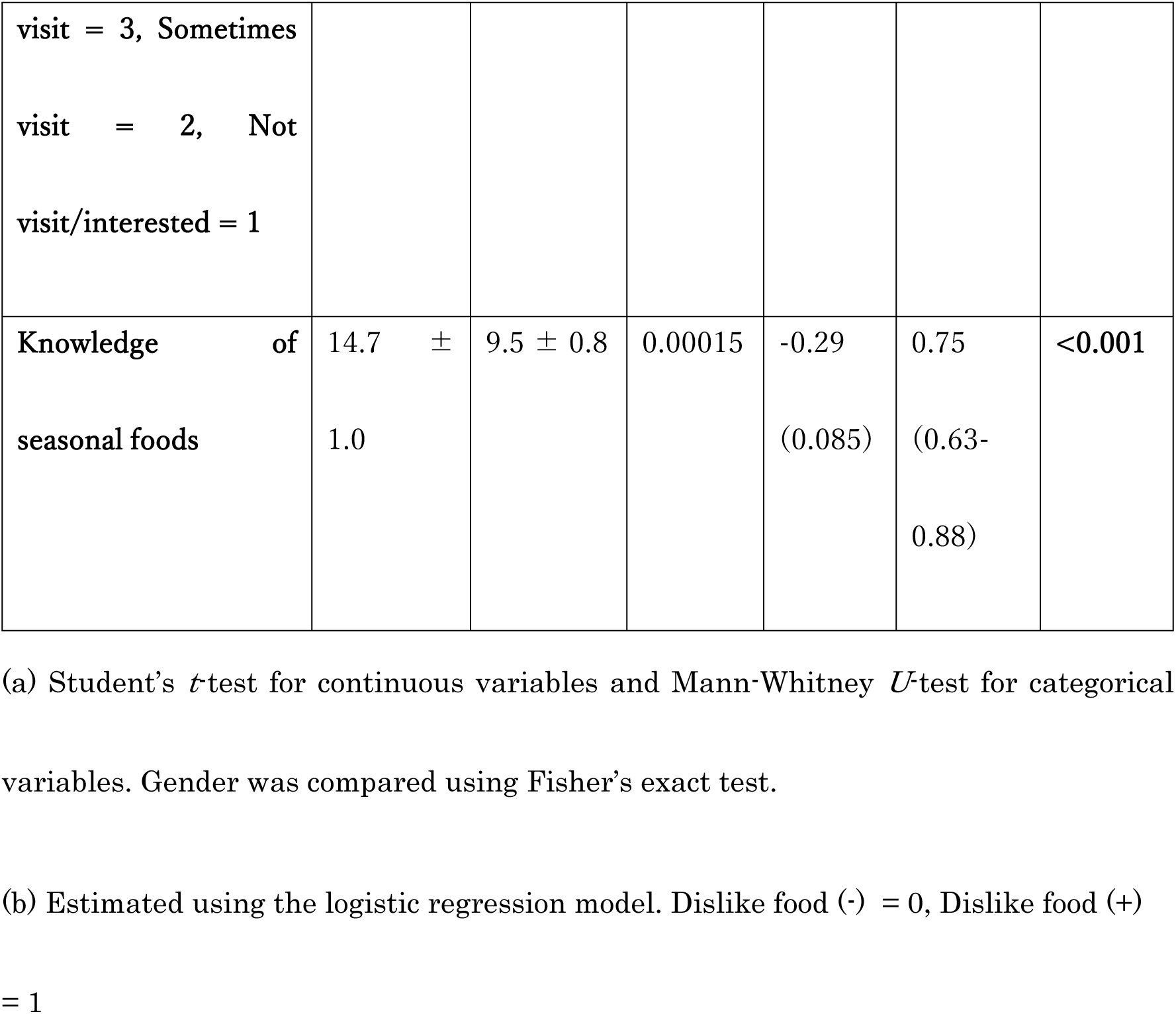
Multiple comparisons of having the dislike food and results of questionnaires.

As shown in Fig. 4, the MPFC response during CD eating tend to high compared with those during eating PD (parried *t*-test. *p* = 0.062). In case of participants who has no dislike food, the MPFC response during CD eating significantly high compared with those during eating PD (*p* = 0.027). In other hands, the MPFC response of the participants who has dislike food during CD eating not significantly different compared with those during eating PD (*p* = 0.58). These changes were not detected in “Just-looking” condition.

**Fig. 4.**
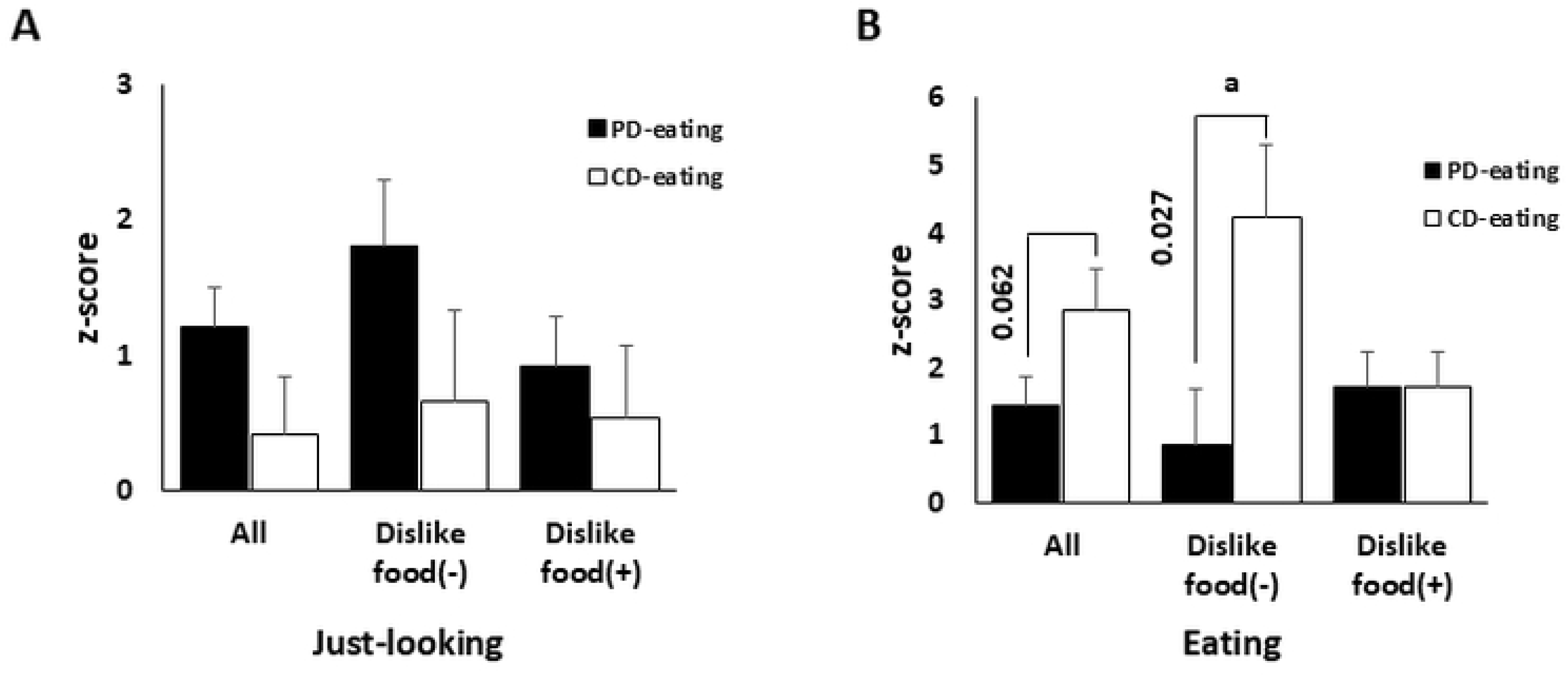
Food preference affects the activity of MPFC during CD and PD eating. (A) Activity of MPFC during “just-looking” when PD eating tend to high compared with those in CD eating. But it was not statistically different. (B) Activity of MPFC during PD-eating was significantly small compared with those in CD-eating in dislike food (-) participants. a indicates *p* < 0.05 by paired *t*-test.

Taken together, we concluded that the activity of MPFC during eating varies depending on the food preference and food intake habits. Moreover, participants who have excellent knowledge of food showed a high activity of MPFC during eating CD (or unknown dishes) but a low activity during PD eating (or known dishes. See also Discussion).

## Discussion

In this study, we found that the activity of MPFC while “just-looking” or “eating” some dishes differed depending on the food preference, food intake frequency, and knowledge of foods. In particular, food preference contributes to the activity of MPFC during eating CD.

As shown in Table 4 and Fig. 4 participants who have no dislike food had more knowledge of seasonal foods and the activity of MPFC during CD eating was high. However, they showed decreased activity of MPFC when they ate PD, which they chose themselves. The activity of MPFC modulates behavior including eating [9] and category-dependent preference signals found in MPFC [14]. The activity of MPFC is also correlated with schematic prior knowledge; it contributes to memory recall or understanding of future events [15,16]. This function could also be associated with eating behavior. Thus, food preference and knowledge of foods affect the activity of MPFC during eating. This study indicated that the activity of MPFC recorded by fNIRS may be used to identify people who prefer provided meals (CD-responders; meals usually provided in hospitals) from those who prefer self-selected meals (PD-responders; such as meals provided in buffet style).

The next step of this study will be performed on elderly participants. However, there are still several concerns. Olfactory, taste, and visual functions decrease with aging [17–19]. For example, the activity of MPFC during an olfactory task was found to decrease with aging [20]. Thus, we should expand the age range of participants such as middle age (around 40 years old) and early elderly age (60 years old without appetite loss) before observing the main target age group (over 80 years old with appetite loss). Moreover, in this study, the participants ate whole dishes, but which are sometimes difficult for elderly people to eat reasons such as the amount of foods is large, swallowing difficulty, and the time course of testing. For example, using specific test foods such as small rice balls, pieces of meat, or food paste could be considered. It was also a problem that the number of responders was small, and there may some possibilities that we under-/overestimated the results of questionnaires even if this study was a pilot study.

## Conclusion

Observing the activity of MPFC during eating by fNIRS potentially enables the separation of the elderly into groups depending on food preference/intake frequency. Further technical improvements will be required to determine whether the food intake test combined with fNIRS will contribute to the treatment of disorders in the elderly such as anorexia.

## Acknowledgements

MYU Research Ltd. proofread this paper.

## Author Contributions

YT planed the experiment, performed the experiment, analysis, and wrote manuscript. YS performed the experiment. TO performed the experiment and wrote manuscript. HS supervise the statistically analysis.

## Data availability statement

We open the data if it requested

## Declarations of interest

none

## Funding sources

This study was supported by Grant-in-Aid for Research Activity Start-up (20K23265) for YT

## Notes

### Competing Interest Statement

NO authors have competing interests

